# Construction and representation of human pangenome graphs

**DOI:** 10.1101/2023.06.02.542089

**Authors:** Francesco Andreace, Pierre Lechat, Yoann Dufresne, Rayan Chikhi

## Abstract

As a single reference genome cannot possibly represent all the variation present across human individuals, pangenome graphs have been introduced to incorporate population diversity within a wide range of genomic analyses. Several data structures have been proposed for representing collections of genomes as pangenomes, in particular graphs. In this work we collect all publicly available high-quality human haplotypes and constructed the largest human pangenome graphs to date, incorporating 52 individuals in addition to two synthetic references (CHM13 and GRCh38). We build variation graphs and de Bruijn graphs of this collection using five of the state-of-the-art tools: `Bifrost`, `mdbg`, `Minigraph`, `Minigraph-Cactus` and `pggb`. We examine differences in the way each of these tools represents variations between input sequences, both in terms of overall graph structure and representation of specific genetic loci. This work sheds light on key differences between pangenome graph representations, informing end-users on how to select the most appropriate graph type for their application.

## 1 Introduction

In recent years, the majority of studies on human genetics have been conducted on the basis of comparing new samples against a single, standard reference sequence. This reference sequence is a linear succession of nucleotides that acts as a blueprint of the human genome. It is routinely used to align raw sequencing data to it in order to find variations between genomes, e.g. single-nucleotide polymorphisms (SNPs), insertions or deletions (indels). It also is the backbone of the UCSC Genome Browser [16] which enables inspection of genomic and epigenomic features. Despite updates that have improved the quality of the human reference sequence in the last two decades, its linear form severely limits the ability to capture population genetic diversity. For instance the locations of frequently observed structural variations cannot be easily integrated into a linear reference. To see this, consider the difficulty of designing a suitable coordinate system in the presence of (possibly nested) structural variants. Having a single genome as reference sequence also introduces an observational bias towards the chosen alleles that were integrated into that sequence, negatively impacting many primary analyses such as reads mapping, variant calling, genotyping and haplotype phasing. As a result our ability to precisely characterize structural variants, SNPs and small indels is limited [5, 12, 30]. The current version of the reference genome (GRCh38) is estimated to miss up to 10% of our species genetic information [28].

Improvements in sequencing data quality and length, as well as genome assembly methods, are providing a fast expanding collection of haplotype-resolved human genome assemblies. If adequately combined together, these high-quality individual genomes may offer an powerful alternative to the linear reference. There now is an active line of research on pangenomes, i.e. data structures that represent a collection of genomic sequences to be analyzed jointly or to form a reference [5, 32]. Pangenome-based approaches have already been successfully applied to short reads mapping and genotyping [8, 30]. These results pave the way for new applications, e.g. genome-wide association studies, where more precise identification of variants can improve the scope of genetic studies in aging, human diseases, and cancer [5, 32].

Several pangenomic data structures have been proposed: multiple sequence alignments, de Bruijn graphs, cyclic and acyclic variation graphs, and haplotype-centric models that use the Burrows-Wheeler transform [5]. Each of these approaches aim to represent a collection of genomic sequences in an efficient way, to store, visualize, and retrieve differences of interest between the considered genomes. Graph-based pangenome data structures, such as the de Bruijn graph and the variation graph, appear so far to be the most advanced in their ability to handle large amounts of input data. They are capable of representing tens to hundreds of human haplotypes simultaneously. Variations graphs use a sequence graph and a list of paths to store input haplotypes, while de Bruijn graphs store all haplotype *k* -mers annotated by their haplotype(s) of origin.

Scaling pangenome graph data structures to store hundreds of genomes is a challenge that requires significant computational resources and engineering efforts. Many software tools have been created, here we briefly describe major ones. Pantools [27] and Bifrost [18] are two methods that have been developed to generate pangenomes for analysis on large collections of genomes, mostly for applications in phylogenetics and bacterial genomics. The PanGenome Graph Builder (`pggb`) [11], `Minigraph-Cactus` and `TwoPaCo` [23] are methods for building general-purpose pangenome graphs. `Minigraph` [21] builds a particular type of pangenome graph by aligning sequences in an iterative way to a reference template. Minimizer-space de Bruijn graphs (`mdbg`) [9] are variants of de Bruijn graphs that can efficiently represent very large collections of bacterial pangenomes (e.g. 600,000 bacteria).

Many human pangenomes have been generated, e.g. using Pantools [27] (7 genomes), `Minigraph` [21] (94 haplotypes), `Minigraph-Cactus` [2, 17] and `pggb` [22] (94 single chromosomes), and `TwoPaCo` [23] (100 simulated genomes). Lastly, a draft version of a human reference pangenome constructed using `pggb` and the `Minigraph-Cactus` pipeline has appeared in a very recent article from the Human Pangenome Reference Consortium [22]. In this article we provide a comprehensive view of whole-genome human pangenomics through the lens of five methods that each implement a different graph data structure: `Bifrost`, `mdbg`, `Minigraph Minigraph-Cactus` and `pggb`. We examine several features of pangenome graphs, in particular their scalability and how they represent genetic diversity. To this end we collected all publicly available high-quality human haplotypes (section 2.1), and attempted to construct pangenomes of various complexity with each selected tool (section 2.2).

The rest of this article is structured as follows. We describe in more details the human haplotypes and pangenome construction tools in the Section 3. In Section 4 we focus specifically on two HLA loci in order to observe fine-grain difference between the methods. In Section 5 we examine other characteristics of the methods such as their ability to perform dynamic updates, remain stable under permutation of inputs, being accessible to downstream applications and ability to visualize the generated graphs. Finally we discuss in Section 6 an overall assessment of the tools and some perspectives.

## 2 Methods

### 2.1 Datasets and haplotypes collection

In order to evaluate the state of current human pangenome representations, we sought to build a human pangenome that contains all publicly available high-quality human haplotypes. We collected from two different sources 102 different haplotypes from the genome of 51 individuals, and also used the two reference genomes, GRCh38 from the Genome Reference Consortium (GRC) [26] and CHM13 v2.0 cell line of the T2T Consortium [24]. Five haplotypes correspond to Google Brain Genomics DeepConsensus [3] assembly dataset: they are hifiasm assemblies of PacBio Hi-Fi reads corrected with DeepConsensus. The average of their N50 is 37,99 Mbp. The remaining haplotype assemblies as well as the T2T reference are from the Human Pangenome Reference Consortium (HPRC) year-1 freeze [32], and GRCh38 is from the GRC. Their average N50 is 40.3 Mbp. Since HG002 is contained in the DeepConsensus data, the HPRC HG002 haplotypes were not used. The origin and the sex of the individuals are diverse to aim for a fair representation of the diversity in human population: out of 51 total individuals, 21 are males and 30 are females and they represent 14 different ethnic groups, from US to Africa and Asia.

To evaluate the scalability of pangenome construction tools, we created three datasets of increasing size: 1) 2 haplotypes from the same individual, HG006, 2) 10 haplotypes from 5 different individuals (HG002, HG003, HG004, HG006 and HG00735) and finally 3) all of the 104 haplotypes. To test whether the order of the input sequences matters, we considered various random orderings for the 10 haplotypes in Dataset 2. Since `Minigraph` needs a reference sequence as fist haplotype in order to correctly build the graph, we generated specific 2 and 10 haplotypes datasets with the first haplotype replaced by the reference genome CHM13. This was applied to the `Minigraph-Cactus` pipeline as well as it uses `Minigraph` variation graphs.

### 2.2 Pangenome graph construction tools

We evaluated tools that generate graph pangenomes as variation graphs and colored compacted de Bruijn graphs. Variation graphs are generally locally acyclic while de Bruijn graphs have cycles. In variation graphs, nodes represent sequences and edges represent immediate sequence adjacency without overlap. Variation graphs are generally easier to visualize and to interpret while challenging to construct at scale and, apart from `pggb`, require a reference genome. In de Bruijn graphs (dBG), nodes are *k* -mers (string of length k) and edges are (k-1)-overlaps between nodes. In practice, dBGs are represented in a compact way where all nodes along unbranching paths are compacted into *unitigs*. The resulting graph is called compacted De Bruijn Graph, where nodes are unitigs and edges represent (k-1)-overlaps. As shown in Figure 1, de Bruijn graphs result in large graphs that pose visualization and interpretation challenges, in particular as there is no alignment to a reference.

**Figure 1.**
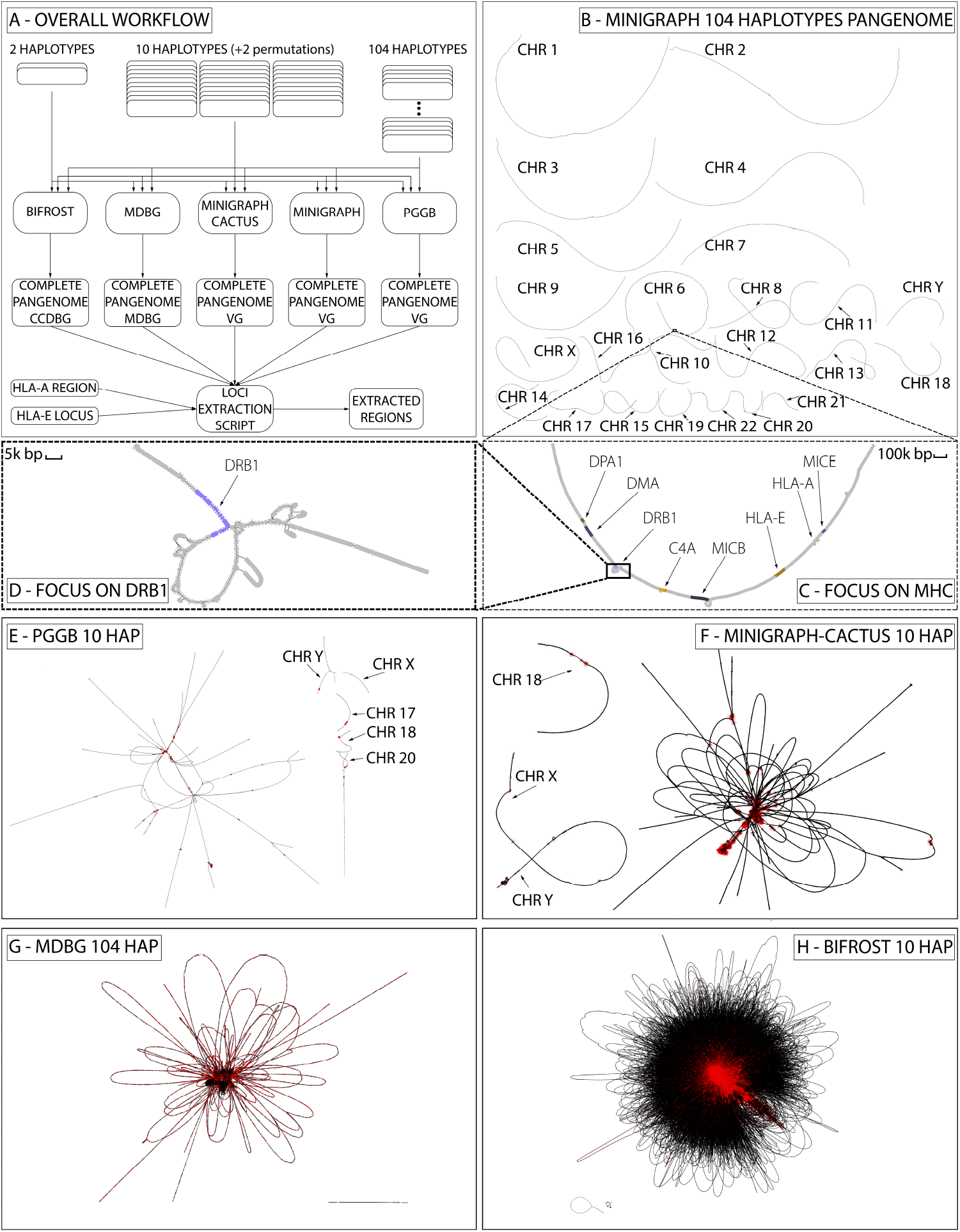
The complete pangenome construction scheme and visualization. **A**, The overall workflow, using 5 different tools on 3 different datasets; **B**, complete 104 haplotypes variation graph built by `Minigraph`; **C**, focus on part of HLA (MHC) region in chromosome 6 from panel B; **D**, focus on DRB1-5 locus of HLA from panel C; **E**, complete 10 haplotypes variation graph built with `pggb`; **F**, 10 haplotypes variation graph built with `Minigraph-Cactus`; **G**, 104 haplotypes pangenome `mdbg`; **H**, 10 haplotypes `Bifrost` dBG. All graphs except those produced by `Minigraph` have been simplified using gfatools and rendered using `Bandage`. VG is for variation graph.

- `Bifrost` constructs dynamic, coloured compacted de Bruijn Graphs (*cdBG*). It first generates a standard dBG using an efficient variant of Bloom Filters and then computes the compacted dBG from it. Colors, i.e. identifiers representing the sample origin of each k-mer are added by storing an array per *k* -mer. A human genome cdBG typically consists of a single large connected component, as common *k* -mers are shared between chromosomes. This pangenome representation contains all the variations present in input sequences.
- `mdbg` builds a variant of de Bruijn graphs called a minimizer-space de Bruijn Graph (`mdbg`), which is efficient to construct as it only considers a small fraction of the input nucleotides. Color information is currently not supported in the implementation. Similarly to Bifrost, a `mdbg` also typically represents a human genome as a single large connected component, albeit with orders of magnitude less nodes. Minimizer-space de Bruijn graphs mostly discard small variants, yet are sensitive to heterozygosity which creates branches in the graph.
- `Minigraph` constructs a directed, bidirected and acyclic variation graph iteratively by mapping new haplotypes using a combination of the minimap2 tool and the graph wavefront alignment algorithm. The first input sequence acts as a backbone for the whole representation. The sample(s) of each node are stored in a rGFA output file. `Minigraph` considers only variations longer than 50 bps hence it is oblivious to isolated SNPs and small indels: even if it produces base-level alignment for contigs, the graphs are not a base-level resolution. The resulting graph is divided into connected components that correspond to the chromosomes present in the first given input genome.
- `Minigraph-Cactus` is a variation graph construction pipeline that combines `Minigraph` to generate a structural variations graph and `Cactus` base aligner to generate base-level pangenome graphs of a set of input assemblies and embed haplotypes paths. `Cactus` [2] is a highly accurate and scalable reference-free multiple whole-genome alignment tool, that in this pipeline considers the reference sequence used by `Minigraph` to ensure that the resulting variation graph is acyclic. The final graph is further normalized using GFAffix [7]. The pipeline allows to generate multiple graphs, one for each chromosome, or produce a single graph that includes inter-chromosomal variants.
- `pggb` is a directed acyclic variation graph construction pipeline rather than a single tool. It calls three different tools: pairwise base-level alignment of haplotypes using wfmash [15], graph construction from the alignments with seqwish [10], graph sorting and normalization with smoothxg and GFAffix [7, 13]. The resulting variation graph represents variations of all lengths present in the input sequences.

## 3 Scalability and characteristics of pangenome graph construction tools

We ran the above five tools on the three datasets of Table 1, and compared their computational performance as well as characteristics of the produced pangenome graphs. The goal is to assess the ability of each method to scale to data available in the near future, i.e. thousands or even millions of human genomes [28].

**Table 1.** Description of the three datasets generated to test the scalability of the tools.

| Haplotypes | Project | Bases |
| --- | --- | --- |
| 2 | Google<sup>1</sup> | 5.9 Gbp |
| 10 | Google, HPRC<sup>2</sup> | 30 Gbp |
| 104 | Google, HPRC, 1KG<sup>3</sup>, T2T<sup>4</sup> | 313.6 Gbp |

**Table 2.** URL, version, pangenome representation and parameters of the three analyzed tools. pggb/0.2.0 used wfmash v0.7.0, seqwish v0.7.3 and smoothxg v0.6.1.

| Tool | Github repository | Graph type | Version | Parameters |
| --- | --- | --- | --- | --- |
| **Bifrost** | pmelsted/Bifrost | De Bruijn graph | 1.0.5 | -k100 -c |
| **pggb** | pangenome/pggb | variation graph | 0.2.0 | -p 98 -s 10000 -k 311 -G 13033,13117<br>-O 0.03 -v -t 8 -T 8 -A -Z |
| **Minigraph** | lh3/Minigraph | variation graph | 0.18 | -cxggs |
| **Minigraph-Cactus** | ComparativeGenomics<br>Toolkit/cactus | variation graph | 2.2.3 | -maxLen 10000 -delFilter 10000000 |
| **mdbg** | ekimb/rust-mdbg | De Bruijn graph | 1.0.1 | -k 10 -d 0.0001 -minabund 1 -reference |

### 3.1 Computational metrics

The performance of each tool is evaluated in terms of running time, peak memory, disk space required by the output data structure (graph and annotations). We also compared the number of nodes, edges and connected components as indicators of the complexity of the graph. Results are displayed in Table 3.

**Table 3.** Time, memory, final disk space, nodes, edges, total connected components and connected components with more than 1% of base pairs comparison of `Bifrost`, `mdbg`, `pggb`, `Minigraph` and `Minigraph-Cactus` for different number of haplotypes in input. `Minigraph-Cactus` times include the `Minigraph` graph construction step. `pggb` was not able to complete its execution on the largest dataset in more than 2 weeks thus it is not considered. `Minigraph-Cactus` failed to compute the 104 HAP dataset.

| Haplotypes | Metric | Bifrost | pggb | Minigraph | Minigraph-Cactus | mdbg |
| --- | --- | --- | --- | --- | --- | --- |
| 2 | time (hh:mm:ss) | 1:21:25 | 15:45:30 | 00:08:33 | 3:11:59 | 00:02:38 |
|  | memory (GB) | 53 | 24 | 38 | 66 | 8 |
|  | disk space (GB) | 4.8 | 4.3 | 2.9 | 3.6 | 4.4 |
|  | nodes | 9,482 k | 8,492 k | 34 k | 10,851 k | 17 k |
|  | edges | 13,108 k | 11,503 k | 48 k | 14,702 k | 23 k |
|  | conn comp | 2 | 1402 | 25 | 4 | 174 |
|  | conn comp > 1% bp | 1 | 30 | 24 | 4 | 24 |
| 10 | time (hh:mm:ss) | 2:27:29 | 117:08:09 | 2:03:29 | 15:57:05 | 00:05:46 |
|  | memory (GB) | 102 | 71 | 49 | 154 | 18 |
|  | disk space (GB) | 12 | 7.6 | 2.9 | 7 | 9.7 |
|  | nodes | 27,468 k | 29,315 k | 133 k | 37,767 k | 67 k |
|  | edges | 37,584 k | 40,282 k | 190 k | 51,595 k | 93 k |
|  | conn comp | 3 | 864 | 25 | 3 | 40 |
|  | conn comp > 1% bp | 1 | 5 | 24 | 3 | 1 |
| 104 | time (hh:mm:ss) | 18:38:28 | — | 46:22:00 | — | 00:31:38 |
|  | memory (GB) | 122 | — | 61 | — | 39 |
|  | disk space (GB) | 29.4 | — | 3.2 | — | 38 |
|  | nodes | 106,339 k | — | 632 k | — | 270 k |
|  | edges | 293,839 k | — | 912 k | — | 396 k |
|  | conn comp | 17 | — | 25 | — | 1097 |
|  | conn comp > 1% bp | 1 | — | 24 | — | 1 |

In terms of running time, `mdbg` is two orders of magnitude faster than other tools on all considered datasets, taking around two minutes on the H2 dataset and half an hour on H104. `Bifrost` is the second fastest on H104 (18 hours), and `Minigraph` is the second fastest on H2 (8 minutes). `Minigraph-Cactus` takes one order of magnitude more time than `Minigraph`. We could not obtain graphs for `pggb` and `Minigraph-Cactus` on H104 as for the first execution did not finish after 2 weeks and the second returns an error.

In terms of memory usage, `mdbg` consistently uses less than half the memory of other tools (31 GB on H104), followed by `Minigraph` (61 GB on H104). On H2 all tools used between 8 and 66 GB of memory.

All tools used reasonable disk space to store the resulting graph, *≤* 12 GB for H10 and *≤* 38 GB for H104. Although `Minigraph-Cactus` and `pggb` retain all variations and are the only two tools able to reconstruct the input haplotypes directly from the graph, they are the second and third most efficient in term of disk space (for `Minigraph-Cactus`, 3.6 GB on H2 and 7 GB on H10). While `Bifrost` and `Minigraph` perform all computation in memory, `pggb`, `Minigraph-Cactus`, and `mdbg` store intermediate files on disk, taking comparable space to the input size (up to 3x for `Minigraph-Cactus`).

### 3.2 Different tools yield different pangenome graphs topologies

Graph metrics such as the number of nodes, edges and connected components provide useful insights on the level of detail of the represented variations and on the complexity and accessibility of the information inside the pangenome.

The number of graph nodes varies between 17,000 and 11 millions for the H2 dataset across all tools. In all cases, the number of nodes is at least 3 orders of magnitude smaller than the number of bases in the haplotypes, indicating that pangenome graphs are effective at compressing linear parts of the haplotypes. Tools which discard variations (`Minigraph` and `mdbg`) yield in the order of 10^4^–10^5^ nodes across all datasets, while tools which retain all variation (`Bifrost`, `Minigraph-Cactus` and `pggb`) yield in the order of 10^6^–10^7^ nodes. In all cases going from the H10 dataset to the H104 dataset increases the number of nodes by 5x, indicating that graph complexity grows sublinearly with the number of added haplotypes.

The number of connected components varies between 2 and 1402 across all methods and datasets, and the number of large components (i.e. those with more than 1% of total base pairs) varies between 1 and 30. If chromosomes were separated perfectly, pangenome graphs should contain exactly 24 connected components (one per nuclear chromosome, excluding mitochondria). `Minigraph` produces 24 large connected components as the number of chromosomes in the reference CHM13 v2.0 (25 including mitochondria). `Bifrost` and `Minigraph-Cactus` yield graphs with less than 25 connected components while `mdbg` and `pggb` have more than 25. In the `Bifrost` dBG, the vast majority of sequences (*>*99.99%) are in a single giant component, as chromosomes are joined because they share common *k* -mers. In `mdbg` such joining does not occur on dataset H2, which has 24 large enough components (each containing *>* 1% of bases), possibly due to the absence of long and similar enough regions between chromosomes. `Minigraph` does not map any mitochondrial sequence from the input haplotypes to the reference, while they do get included into `Minigraph-Cactus` graphs.

Even if it is common practice to analyze pangenomes chromosome by chromosome [17, 22], in this analysis we purposely used entire genomes as input instead. This was done for two reasons: i) to highlight the scalability of the tools, and ii) because separating chromosomes prevents the identification of inter-chromosomal inversions, translocations, and transposable elements, even if most of the generated inter-chromosomal events are probably alignment artifacts. The effects of this choice can be seen in the `pggb` and the `Minigraph-Cactus` H10 variation graphs of Figure 1. In the `pggb` graph 19 chromosomes are linked into a single giant component, while chromosomes 17, 18, 20, X, and Y are in other large components. This giant component consists of 25 M nodes that contain 83% of the total basepairs. The remaining 859 components represent only 4.7% of the total bases due to small sequences in the input haplotypes. In the `Minigraph-Cactus` graph all chromosomes are linked into a single giant component except chromosome 18 that is in a separate component, and the sexual chromosomes (X and Y) that are connected together into another component.

## 4 Interpretation of variation in pangenome graphs: focus on two HLA loci

The ability to detect and annotate variations among input haplotypes defines the scope of each pangenome graph construction method. Previous work [4] recommends to build graphs on a specific loci rather than the entire genome for the purpose of i) identifying genomic diversity and ii) mapping raw reads to divergent regions, specifically difficult-to-map repeats. Here we evaluate how pangenomes built from entire haplotypes represent specific biologically relevant loci.

### 4.1 Extraction of HLA-E and a complex HLA region from complete pangenome graphs

We extracted from complete pangenomes the regions corresponding to two loci of the Human Leukocyte Antigen complex, also known as HLA. These regions are highly medically relevant as they contain many disease-associated variants [6]. The first locus is the HLA-E gene, that is part of the nonclassical class I region genes, spanning 4,8 kbp and is relatively conserved across populations. It has been shown to have significant association with hospitalization and ICU admission as a result of COVID-19 infection [31]. The second is an HLA complex region comprising the HLA-A gene, part of the classical, highly polymorphic class I region. It is around 58 kbp long and contains the HLA-U, HLA-K, HLA-H, and HCG4B genes. We extracted these two regions from pangenome graphs using a custom script that yields a subgraph corresponding to a given set of sequences and their variation. The script uses a different recommended method for each of the pangenome graph types. In a nutshell, we extracted regions using exact coordinates when possible, and and resorted to sequence-to-graph alignment otherwise (see Appendix section 7.2 for details). While on variation graphs and mDBGs nearby nodes of an aligned region correspond to variations of the locus, this is not always true for standard dBGs. Extracting accurate and complete loci representation is an unsolved challenge for dBGs.

### 4.2 HLA-E: a low complexity region

Figure 2 shows how the different tools represent HLA-E over datasets H2, H10 and H104. As expected, `Minigraph` does not detect any variation, since the SNPs that characterize the region are too small to be considered in the construction steps of their algorithm. `pggb`, on the contrary, has 2 SNPs in H2 and 3 in H10. `Bifrost` detects the same SNPs as `pggb` in H2 and H10. Both of them represent the exact same variations and render the same haplotypes paths. `mdbg` captures the heterozygosity of a large region containing the HLA-E locus as the number of samples grows. As the `mdbg` graph is built in minimizer space, nodes represent long genomic segments (in the order of hundreds of thousand of base-pairs). In H10 and H104,the minimizer-space representations of the haplotypes are identical; however differences in flanking regions of the graph create variations that are captured in extra nodes that are also extracted in this region. On H2, `Minigraph-Cactus` detects 3 variations as the dataset used is different, containing the CHM13 reference and just one haplotype of HG006 (as in `Minigraph`), as discussed in Section2.1.

**Figure 2.**
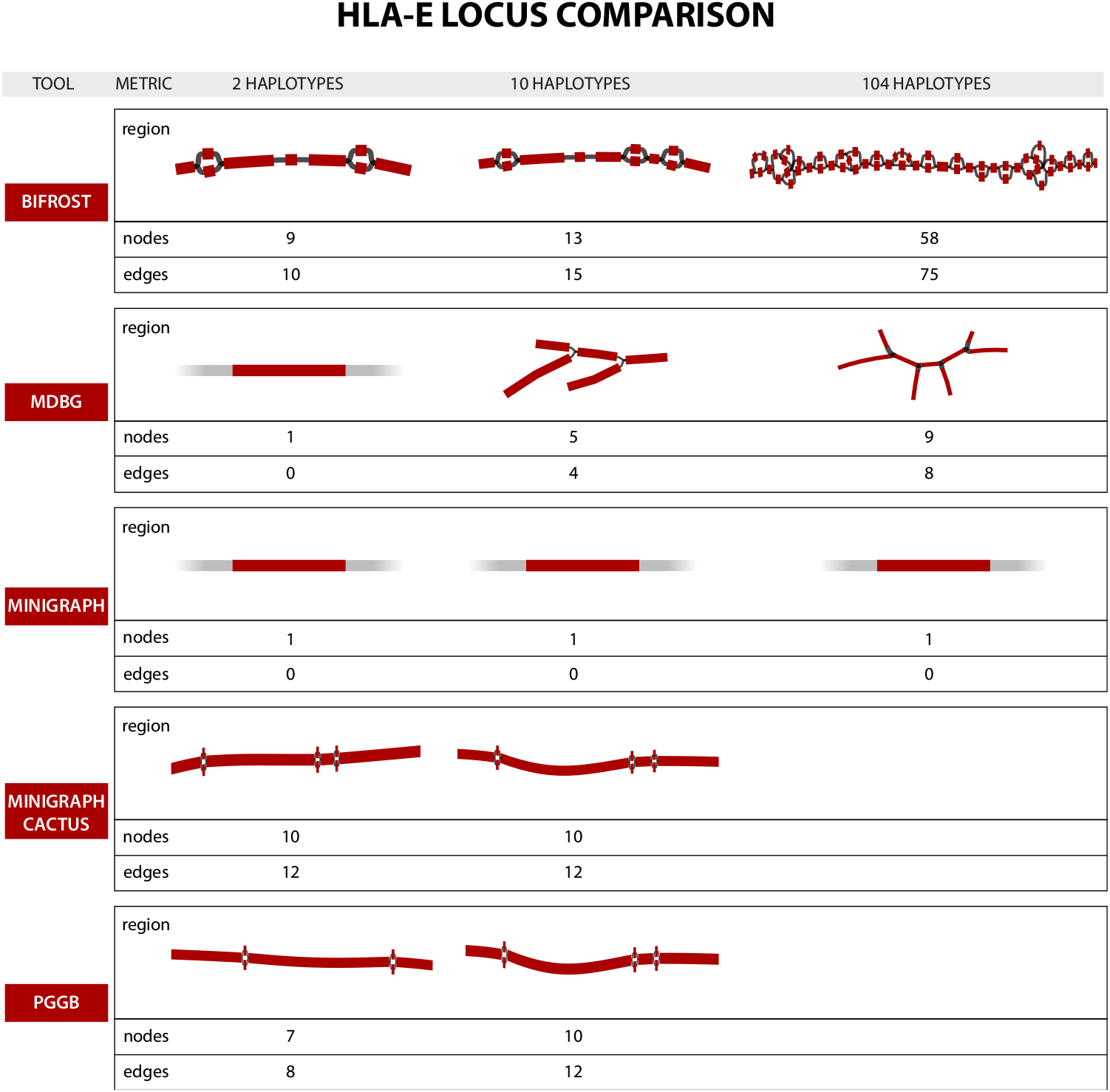
Representations of the HLA-E locus by five graph construction methods over three increasing large human pangenomes. Nodes highlighted in red contain part of the locus sequence. The numbers of nodes and edges displayed below each graph concerns the whole subgraph (both red and grey nodes). `Minigraph`, on H2, H10 and H104, and `mdbg`, on H2, have only a portion of one node highlighted since the 4.8k bp region is contained inside a single, long node.

Figure 2 also illustrates how pangenome complexity grows with the number of genomes: the `Bifrost` H104 subgraph has the most variation across all methods, highlighting that dBGs represent variations exhaustively in large graphs. On the other hand, `pggb` has the most straightforward method for extracting subgraphs, and also represents variants exhaustively in datasets H2 and H10, but could not scale to the H104 dataset.

### 4.3 HLA complex locus: high complexity region

Figure 3 is the counterpart of Figure 2 for the complex locus part. In this case the overall interpretability of the region is more challenging, as the number and the structure of the variations is different than in HLA-E. It is also more difficult to compare across tools. Base-level variations, e.g. SNPs, are not visually recognizable in Figure 3 in the methods that retain them (i.e. `pggb`, `Minigraph-Cactus` and `Bifrost`) due to the large sizes of graphs.

**Figure 3.**
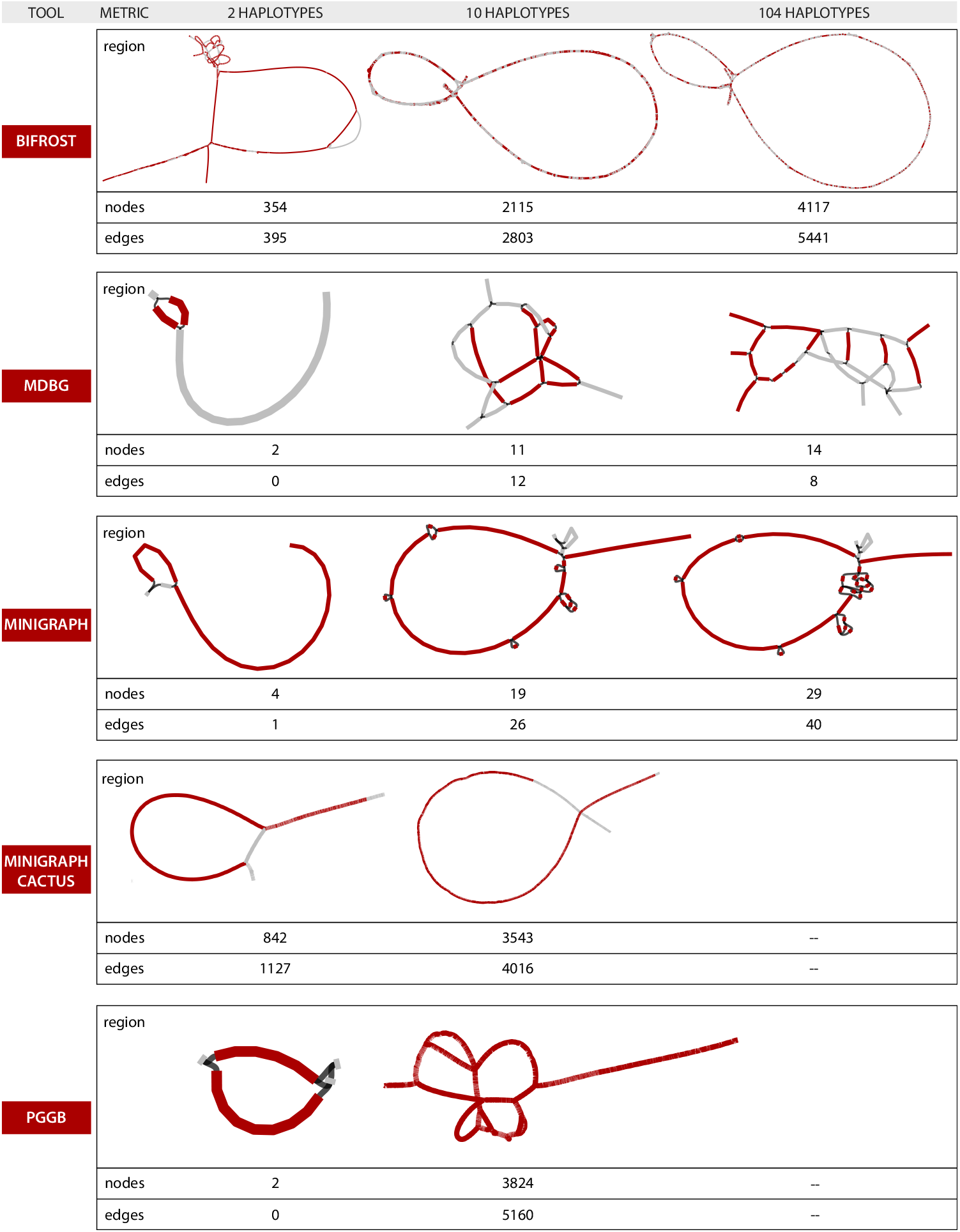
Representations of the complex HLA region by five graph construction methods over three increasing large human pangenomes. See caption of Fig. 2 for details.

There are notable differences in how tools represent the variation, which is well-illustrated in the H2 dataset. While `Minigraph` renders H2 as a single sequence plus a large structural variant (SV) of *≈* 52k bp, `pggb` separates it into two paths that differ by *≈* 54k bp in length. `Bifrost` represents a detailed bubble that contains many variations inside each path. In `mdbg`, even extracting the complete locus is a challenge as many of the subgraph nodes were not selected by our procedure. `Minigraph-Cactus` adds base level divergences between haplotypes on top of `Minigraph` SV graph.

These differences between representations are further accentuated in the H10 dataset. For it, `pggb` tends to separate the haplotypes into different paths, `Bifrost` renders consistently the same compacted representation and `Minigraph` neglects most of the small differences but is able to display accurately the general picture, and `Minigraph-Cactus`, as in H2, adds small variations on top of `Minigraph` structure.

## 5 Uncovering characteristics of graphical pangenome tools

The data structures generated by pangenome building tools are expected to facilitate comparisons between the input genomes. In addition pangenome graphs should be stored in such a way to be easily used by downstream applications. We identify 8 important features for pangenome graph construction tools: i) stability, ii) editability, iii) accessibility by downstream applications, iv) haplotype compression performance v) ease of visualization, vi) quality of metadata and annotation. Two other but important features, scalability and interpretability of produced graphs, were already discussed in Sections 3 and 4. Table 4 summarizes some of the following considerations on the relative strength of the tools.

**Table 4.**
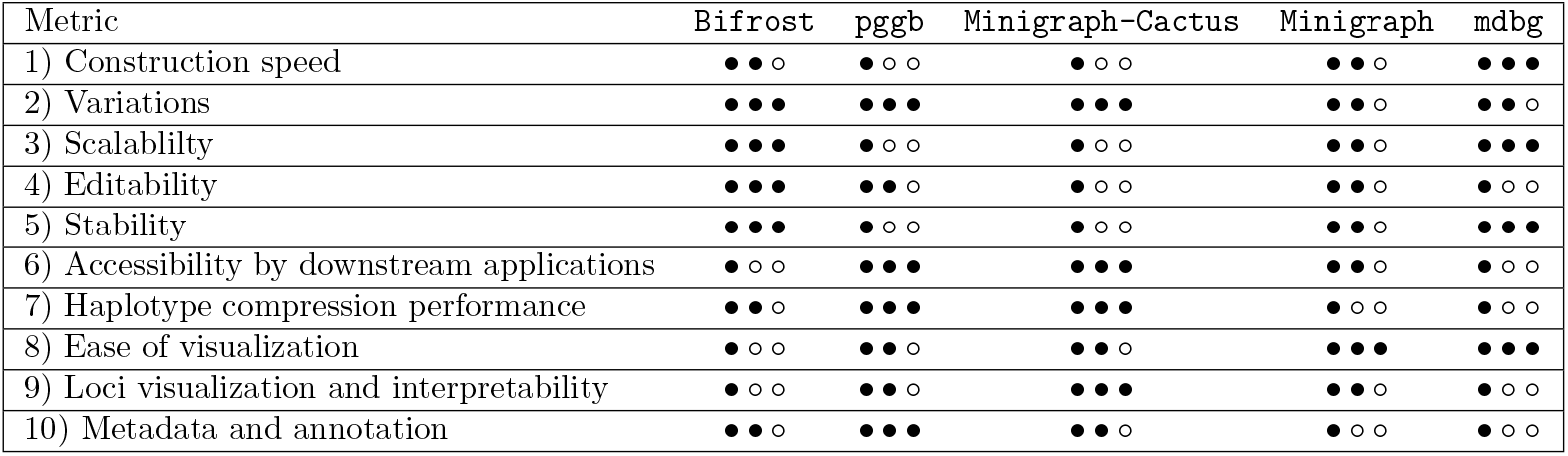
Relative strengths of five pangenome graph construction tools. Explanation of rows: 1) Efficacy of construction algorithm, measuring wall-clock time; 2) degree to which variants (e.g. SNPs) are retained; 3) ability of a tool to perform well on large datasets, both in comparison to other tools but also compared to smaller datasets; 4) ability to modify the produced data structure to add or remove haplotypes; 5) property of producing the same result irrespective of perturbations, such as permutation of the input order, and repeating the same run; 6) existence of tools (and operations) that can be applied to the resulting graphs; 7) whether input haplotypes information is retained by the tools, and if so, its space efficiency; 8) whether the entire graph can be directly visualized and interpreted; 9) easiness of ‘zooming in’ a specific genomic region and interpret variants; 10) summarizes the functionalities provided by the tools to annotate the pangenomes with genomic data.

### 5.1 Editability and dynamic updates

As more high quality assemblies will be generated in the near future, haplotypes may be added to a pangenome, or replaced by improved versions. Updating an existing data structure instead of rebuilding it from scratch is both computationally and energetically efficient. However, many succinct data structures currently used in pangenome representation are static, i.e. cannot be updated. Some methods allow a restricted set of editing operations. `Minigraph` allows to add new haplotypes on top of an already built graph. `Bifrost` provides C++ APIs to add or remove (sub-)sequences, *k* -mers and colors from the ccdBG. `pggb`, using `odgi` [14], allows specific operations that delete and modify nodes and edges and add and modify paths through the graph. As `Minigraph-Cactus` can be opened with `odgi`, it supports the same operations as `pggb`. The current `mdbg` implementation uses a dynamic hash table, but does not expose an interface that supports updates.

### 5.2 Stability

Counter-intuitively, a pangenome graph construction tool may in some cases generate different outputs when executed multiple times with the same haplotypes as input. This *unstability* could be due to a permutation in the order of the sequences given as input, or non-determinism in the construction algorithm. Yet in order to facilitate the reproducibility of results, pangenome building tools should generate an unchanged output from the same set of input sequences, independently of the particular run or the order in which these are given. We performed two tests to evaluate tool stability: i) we run the tools 3 times using as input the same H10 dataset and ii) we run the tools twice on shuffled input sequences, i.e. changing the order of the haplotypes of H10.

`Bifrost` and `mdbg` constructed exactly the same pangenome on every test, as by definition, de Bruijn graphs are stable. `Minigraph` generates identical graphs on identical inputs, but generates slightly different graphs when the input is permuted. Indeed the construction algorithm of `Minigraph` is order-sensitive as it augments the existing graph structure by aligning the next given haplotype to it and adding divergent sequences. `Minigraph-Cactus` generates slightly different graphs on identical input. `pggb` generated slightly different graphs while maintaining the same haplotype sequences in the paths. The overall representation of the input genomes is therefore preserved, while the topology of the variation graph varies. The first two of the three phases of the `pggb` pipeline (all-vs-all alignment and graph imputation) produce the same result on different runs with the same input but differences arise when the order of the input haplotypes changes. Most of the differences in the graph topology are thus due to the final smoothing steps.

### 5.3 Accessibility by downstream applications

To facililate their adoption, pangenome representations should be easily processed by downstream analyses. De Bruijn graphs are challenging to analyze due to their high number of nodes, edges, and redundancy (the *k −* 1-overlaps between nodes). Though De Bruijn graph representations usually support queries of presence/absence on nodes (as in `Bifrost`), they lack tools able to perform more elaborate analyses such as those discussed in Section 4, e.g. incorporating haplotype information at the *k* -mer level. On the other hand, variations graphs with paths provide more flexibility, i.e. as in `pggb` and `Minigraph-Cactus` with the `odgi` visualization toolkit. Finally in `Minigraph`, which considers a narrower spectrum of variants, the absence of path information prevents haplotype-level analysis; haplotypes would need to be manually mapped back to the graph.

### 5.4 Haplotype compression

Building a graph pangenome can be seen also as a way to store, compact and retrieve the input haplotypes. As the number of new assemblies is growing faster than the data storing capacity, pangenomes can potentially help save storage space. This is highlighted by the disk space reported in Table 3, which is consistently smaller than the sum of haplotype sizes for all methods and datasets.

In order to losslessly retrieve the input genomes from a pangenome, the representation has to store all variations from the original haplotype sequences as paths in the graph. `pggb` and `Minigraph-Cactus` fall into this category while the other three considered tools do not store paths, or do not consider all variations, thus they are lossy.

Of note, the GBZ tool [29] enables graph pangenomes that store paths in the GFA file format to be stored in a lossless compressed form. It uses a Graph Burrows-Wheeler transformation to compress the graph in a more efficient way than using gzip [29]. Using GBZ, the pangenomes generated by `pggb` and `Minigraph-Cactus` are losslessly compressed with space gains of 3.5-5x.

### 5.5 Ease of Visualization

Visualizing large graphs which exceed hundreds of thousands of node is a challenge that exceeds the scope of pangenomics. The H104 pangenomes are difficult to visualize. Among the visualization tools considered by the Human Pangenome Reference consortium [32], only `Bandage` is able to display the `Minigraph` or `mdbg` H104 graphs, which contains a few million nodes. We reduced the number of nodes and edges of `pggb`, `Minigraph-Cactus` and `Bifrost` H10 graphs by collapsing isolated subgraphs representing SNPs or indels up to 10k bp (using the command `gfatools asm -b 10000 -u`).

### 5.6 Quality of Metadata and Annotation

Augmenting pangenome structures with information from other omics data would increase pangenome relevance in biological discoveries. As biobanks are rapidly growing, more data is available on regulatory regions, transcriptomics, CNVs and other medically relevant traits [1, 19]. Pangenome data structures could leverage such information, and some of the considered tools offer basic functionality in this sense. `Bifrost` provides a function to link data to graph vertices through C++ APIs. `pggb` and `Minigraph-Cactus`, using `odgi`, offer annotation capabilities through insertion of paths or BED records. `Minigraph` and `mdbg` do not offer any annotation feature.

## 6 Discussion

Five state-of-the-art pangenome graphs construction tools were compared on the representation of up to 104 human haplotypes. The approaches significantly differ in terms of speed, graph size, and representation of variations. We find that it remains computationally prohibitive to generate human pangenome graphs for hundreds of haplotypes, especially while retaining all variations. Each approach has its own set of strengths, and ultimately the choice of the method depends on the downstream application. In addition, several takeaway points emerged from our analysis.

First, our focused analysis of HLA loci revealed that de Bruijn graphs and variation graphs represent genomic variations equally well as pangenomes. While de Bruijn graphs are faster to construct, more stable, and scale better in terms of input size, the resulting graphs are challenging to interpret downstream. Variations graphs on the other hand are more practical to analyze at the expense of a less efficient construction step. Their visualization are more straightforward to interpret, mostly due to not having cycles, and provide insights into loci differences.

Second, we can highlight two categories of pangenomic methods that have distinct application domains. `pggb`, `Minigraph-Cactus` and `Bifrost` store all possible variations, and keep haplotype information as paths or colors. They provide a complete picture of the set of variations in the input genomes which makes them difficult to analyze. `Minigraph` and `mdbg` generate ‘sketched’ pangenome graphs that consider only large variants, ignoring smaller differences, and are more efficient to construct and visualize.

Third, every tool possesses an exclusive set of features. `pggb` facilitates downstream analyses using the companion tool `odgi`. It allows to precisely extract and browse any locus of interest. It is the only tool that generates variation graphs without a reference. It also keeps a lossless representation of the input sequences. `Minigraph` generates a pangenome graph based on a reference sequence taken as a backbone. Its shines in the representation of complex structural variations, but does not include small or inter-chromosomal variations. The pipeline `Minigraph-Cactus`, that uses the `Cactus` base aligner, can be used to add small level variations on top of the `Minigraph` graph and to keep a lossless representation of the input sequences. `Bifrost` illustrates that classical de Bruijn graphs are scalable, stable, dynamic, and store all variations. However, extracting information from them remain a challenge. Lastly, `mdbg` is the fastest construction method which generates an approximate representation of differences between haplotypes.

In our view, future directions for human pangenomes building tools should focus on tackling efficiency bottlenecks, aiming to represent hundreds to thousands of haplotypes. Representations should further be lossless and represent the input haplotypes as paths in the graph. Such features would unlock many other applications such as lossless compression of haplotypes and cancer copy number variant analysis.

Finally, we recognize the need for more user-friendly tools that can be used by biologists and that can translate complicated questions into graph queries. While `odgi` begins to address these questions in variation graphs, other approaches have not yet provided user-friendly interfaces. A package similar to `odgi` for de Bruijn graphs would help fully realize their potential.

## 7 Benchmark infrastructure

Running time of pangenome construction tools was measured as wall clock time and peak memory as maximum resident set size using the `time` command. Other metrics were obtained with custom Python scripts. All benchmarks were performed on a Supermicro Superserver SYS-2049U-TR4, with 3 TB RAM and 4 Intel SKL 6132 14-cores @ 2.6 GHz, using 8 cores.

## Availability of data and materials

The scripts used to generate and analyse the pangenomes can be found at https://github.com/frankandreace/CRHPG. Google Brain Genomic assemblies can be found at https://console.cloud.google.com/storage/browser/brain-genomics-public/research/deepconsensus/publication/analysis/genome_assembly.

HPRC assemblies, CHM13 and GRCh38 can be found at https://s3-us-west-2.amazonaws.com/human-pangenomics/index.html?prefix=working/.

## Author’s contributions

FA, YD and RC conceived and designed the project. FA implemented the scripts. FA and PL ran the experiments. FA, YD, PL and RC wrote the paper. The authors read and approved the final manuscript.

## Supplementary material

### 7.1 Twopaco

We did not consider `TwoPaCo` as it is redundant with `Bifrost`. Both methods construct the same de Bruijn graphs. `TwoPaCo` is a method for constructing ccdBG by finding junction *k* -mers at the boundaries of unitigs or in branching nodes. It consists of two main steps in which it approximates the dBG with a Bloom filter in order to reduce the size of the problem and then runs a two pass highly parallel algorithm on it. It constructs ccdBGs similar to Bifros ones. `Bifrost` is faster, supports edit operations, and accepts also reads other than assemblies as input. We tested both tools on NCBI datasets from three different known human variation regions part of the human leukocyte antigen (HLA) complex: HLA-A, MICB and TAP1. These loci have different number of sequences and have complexity and length. The resulting graphs have exactly the same k-mer content and substantially equal topology. The difference is that `TwoPaCo` considers sequences with IUPAC ‘N’ bases while `Bifrost` does not and that in some cases `TwoPaCo` renders some unitigs split in two or more consecutive nodes.

### 7.2 Loci extraction method

For `Bifrost` and `mdbg` graphs, nodes corresponding to the input sequences are identified with `GraphAligner` [25] and the subgraph is extracted using the `Bandage` *reduce* function. As the aligned nodes are not expected to represent the full diversity of the population in the pangenomes, the considered portion of the graph contains also nodes up to a certain distance from the aligned ones: 2 for `mdbg` and 3 for `Bifrost`. This is an arbitrary number based on the size of the sequences spelled by the nodes and on the considered variations. Artifacts, mostly tips, that are not part of the locus of interest are removed from Bifrost graphs with a custom python script. For `Minigraph` generated graphs, the `Minigraph` own alignment function has been used to identify the nodes and then `Bandage` is used to extract the subgraph. For `pggb`, first we generate a bed file of the position of the region of interest in every haplotype used to construct the graph. The ranges are derived from aligning them to the locus sequence(s) using minimap2 [20]. The graph corresponding to the region is then extracted using the `odgi` extract and `odgi` view functions. For `Minigraph-Cactus` we use the same strategy as `pggb`, with the difference that the bed file is only for the reference CHM13, present in the graph.

On a second round, the annotation of specific loci is accomplished by flagging the nodes where the sequences of the locus/loci of interest align with the subgraph. This allows for the highlighting of different sections within the subgraph, such as genes and regulatory regions of interest.

